# Commentary on the nomenclatural status of *Benthoctopus* Grimpe, 1921 (Cephalopoda: Octopoda) and redescription of “*Vulcanoctopus*” *pseudonymus* (Grimpe, 1922)

**DOI:** 10.64898/2026.11.10.765598

**Authors:** Steve O’Shea

## Abstract

The nomenclatural status of the two octopus genera *Bathypolypus* Grimpe, 1921 and *Benthoctopus* Grimpe, 1921 has been the subject of considerable, at times contentious debate since both genera were first proposed. Of these two genera, *Bathypolypus* has precedence, with *Benthoctopus* relegated to synonymy of *Bathypolypus* in 2002. However, the case for this synonymy was not compelling. The systematic complications generated by this act were subsequently aggravated by the description of another genus, *Muusoctopus* Gleadall, 2004—a name proposed for taxa formerly attributed to *Benthoctopus*—and by molecular data that subsequently demonstrated *Muusoctopus* to be a junior synonym of *Vulcanoctopus* González & Guerra, 1998. This created an untenable situation where multiple competing names were available to apply to many generally deep-dwelling octopus species with two rows of suckers, no ink sac, and either smooth or papillose bodies. Remnants of one important specimen historically attributed to the type species of the genus *Benthoctopus* (*Be. piscatorum* (Verrill, 1879)) are herein redescribed. Historical descriptions of this specimen are combined with modern observations and critically compared with what is reported for other North Atlantic *Bathypolypus* and *Vulcanoctopus* taxa to critique the original synonymy of *Benthoctopus* with *Bathypolypus*. The analysis demonstrates that the original synonymy of *Benthoctopus* with *Bathypolypus* cannot be upheld based on existing data, and the widespread nomenclatural repercussions of this status change are discussed. Finally, the type and only known specimen of “*Vulcanoctopus*” *pseudonymus* (Grimpe, 1922) comb. nov. is partly redescribed to facilitate a broader-scale revision of these problematic North Atlantic octopuses.

## INTRODUCTION

In his revision of octopuses in the genera *Bathypolypus* Grimpe, 1921 and *Benthoctopus* Grimpe, 1921 of the North Atlantic, Muus (2002) concluded that the type specimen of *Be. piscatorum* (Verrill, 1879), the type species of the genus *Benthoctopus*, was an “aberrant” individual of the earlier-described *Bathypolypus bairdii* (Verrill, 1873). To preserve nomenclatural stability, Muus (2002: 204) advocated for designating a new type species for the genus *Benthoctopus*, for he considered *Bathypolypus* to take precedence over *Benthoctopus*. At this time, the names of both *Benthoctopus* and *Bathypolypus* were well-established in the literature, and scientists had some understanding of what each represented—both lacked an ink sac and had two rows of suckers, but *Benthoctopus* species were generally smooth bodied, had a crop diverticulum, a W-shaped funnel organ, and a non-laminate ligula, while those of *Bathypolypus* were generally papillose, lacked a crop diverticulum, had a V V-shaped funnel organ, and their ligula was laminate. Exceptions were known.

In response to Muus’s revision, Gleadall (2004) proposed *Muusoctopus* Gleadall, 2004 as a new genus to accommodate at least *O. januarii* Hoyle, 1885, which had formerly been attributed to *Benthoctopus*. At the time Gleadall (2004: 108) maintained that “the relationship between *Muusoctopus januarii* and other nominal species without an ink sac, previously identified as species of ‘*Benthoctopus*,’ must await thorough revision and redescriptions of other such octopodid species living in deeper waters that currently have not been identified as belonging to the genus *Bathypolypus*”. Although many new species have since been attributed to the genus *Muusoctopus*, there has been no comprehensive reassessment of the work of Muus upon which the original synonymy of *Benthoctopus* with *Bathypolypus* was based, and evaluations of the diversity of “inkless octopus” taxa have remained largely regional.

Gleadall’s rationale for describing *Muusoctopus* was that retaining *Benthoctopus* as a genus distinct from *Bathypolypus* would confer no nomenclatural stability. He considered that its constituent species were “almost certainly polyphyletic” (a sentiment expressed earlier by Norman et al. (1997), and later by Norman & Hochberg (2005) and Vecchione et al. (2009)); that “the genus assignations of a number of species formerly identified to *Benthoctopus* required revision”; and that “nothing was to be gained from reassigning type species for one or more of the genera now synonymized under *Bathypolypus*”. Gleadall (2004) also regarded *Bathypolypus* as having precedence over *Atlantoctopus* Grimpe, 1921.

Subsequent molecular analyses challenged the suggestion that taxa historically attributed to *Benthoctopus* represented a polyphyletic assemblage (Jorgensen et al., 2010; Ibáñez et al., 2016, 2020; Sanchez et al., 2018; Strugnell et al., 2009, 2011). Further, these analyses revealed that another genus, *Vulcanoctopus* González & Guerra, 1998 (in González et al., 1998)—initially considered to be endemic to hydrothermal vents—was nested among taxa attributed to *Muusoctopus*. Consequently, if these taxa are treated as congeneric, *Vulcanoctopus*, having nomenclatural priority over *Muusoctopus*, is the appropriate name to apply to them (O’Shea et al., 2026).

Problems with the taxonomy of these octopuses were recognized long before Muus completed his review. In fact, they began when Grimpe (1921) named both *Bathypolypus* and *Benthoctopus* in a preliminary note without generic diagnoses, in which he designated the poorly described *O. piscatorum* Verrill as the type species of the latter. Robson, who toiled on this group for the ensuing decade, expressed an increased difficulty with upholding a distinction between them in what would be his final contribution on the group (Robson, 1932: 51, 249; O’Shea et al., 2025). Wirz (1955: 146) reached a similar conclusion. However, while Robson and Wirz were questioning whether sufficient differences existed to warrant recognizing two genera, subsequent authors almost universally continued to recognize both genera, but debated whether the type specimen of the type species of each genus was consistent with what these genera had come to represent (e.g., Voss, 1988; Voss in Voss & Pearcy, 1990: 67; Muus, 2002; Gleadall, 2004).

The same molecular analyses that identified monophyly in taxa historically attributed to *Benthoctopus* also clearly demonstrated the existence of two genera (the second typically referred to as *Bathypolypus*). Surprisingly, however, given the historical difficulties in clearly differentiating these two genera, they proved not to be closely related. They were, in fact, accommodated in different families (Strugnell et al., 2014)—the Enteroctopodidae Strugnell, Norman, Vecchione, Guzik & Allcock, 2014 and Bathypolypodidae Robson, 1929—and to complicate matters, molecular data were deemed necessary to correctly assign them. This unexpected result called into question the reliability of those characters used in the generic diagnoses of Thiele (1935) and Voss (1988), upon which taxonomic decisions had been based for almost 85 years. Given the apparent monophyly of this group, pre-molecular-era taxonomists must have used something tangible to consciously attribute species to *Benthoctopus*. However, precisely what morphological criteria these were has yet to be clearly articulated.

Although the most appropriate name to use for species formerly attributed to *Benthoctopus* was recently addressed (O’Shea et al., 2026), more problems exist that could, if not resolved, destabilize the nomenclature of these animals further. One of these is the identity of a large, smooth-bodied octopus with two rows of suckers and no ink sac that Aldrich & Lu (1968) described from Placentia Bay, Newfoundland, and attributed to Verrill’s *Be. piscatorum*. Because Muus (2002) maintained that only one “inkless octopus” (*sensu* Gleadall et al., 2010) species (*Ba. bairdii*) occurred in these waters, evidence for the presence of a second species would require his synonymy of *Be. piscatorum* with *Ba. bairdii* to be reconsidered. If Muus’s identification was incorrect then the genus *Benthoctopus* might remain valid, with ripple effects affecting any nomenclatural act based on his synonymy since. This Placentia Bay specimen is herein redescribed, and how its identity (which, regrettably, cannot be resolved herein) might affect the nomenclature of these octopuses is discussed. This redescription and appraisal are augmented with a (partial) redescription of the type and only known specimen of another problematic inkless octopus taxon from the North Atlantic—the poorly known “*Vulcanoctopus*” *pseudonymus* (Grimpe, 1922). Measurements and counts of both of these species are compared with those for the lectotype of *V. januarii* (Hoyle, 1885)—the type species of *Muusoctopus*—and holotype of *Be. piscatorum*.

## MATERIALS AND METHODS

Definitions of measurements and indices (below) are mostly as detailed in O’Shea (1999) and O’Shea et al. (2026); definitions of counts or indices marked with an asterisk (*) depart notably from those of Roper & Voss (1983). The term “buccal bulb” is used as opposed to “buccal mass”, in accordance with terminology widely applied in other molluscan taxa (e.g., Brenzinger et al., 2013; Jensen, 1996; Rudman, 1984). Descriptions of the beaks and relative sizes of the buccal bulb and posterior salivary glands specifically follow O’Shea (1999: fig. 3, tables 2–4). Because of specimen incompleteness, and because dissection was not permitted (only non-invasive examination was permissible), characters and their states cannot be consistently reported for each specimen, limiting species’ descriptions and what comparisons can be made between them.

Upper case letters (i.e., Table, Figure) are used when referring to tables and figures throughout this text, but when referring to tables, figures, and plates in the works of others, lower case letters (i.e., table, figure (fig.) and plate (pl.)) are used. Synonymies are limited to salient accounts, mostly in which relevant specimens, primarily types, have been examined, described, and/or critically discussed. Citations in checklists are specifically excluded.

### Institutional Abbreviations

Natural History Museum (BM), United States National Museum, Smithsonian Institution (USNM), and Musée océanographique de Monaco, France (MoM).

### Morphological abbreviations

abbreviations in the text and tables (and how they are measured) include: AL—arm length, herein denoted I (dorsal) to IV (ventral), measured from the beaks to the outstretched arm tip; ALI—arm length as a percentage of mantle length (or total length, where specified); ASC—*whole arm sucker count; ASn—normal (not abruptly enlarged) arm sucker diameter; BBL—buccal bulb length (greatest dimension); EO—diameter of the eye orifice; FuL—funnel length (measured from the anterior funnel opening to the posterior border along the ventral midline); FFuL—free funnel length (measured from the anterior opening of the funnel to its point of dorsal attachment to the head); FFuLI—*free funnel length as a percentage of FuL; GiLC—gill lamellae count, inner/outer demibranch; HW—head width (greatest width of head at the level of the eyes); ML—dorsal mantle length (measured from the midpoint between the eyes to the posterior end of the mantle); MW—mantle width (greatest dorsal straight-line width of the mantle); TL—total length of the body (measured from the tip of the mantle to the outstretched arm tips, along the antero-posterior axis); WD—web depth (measured from the beaks to web margin at the mid-point between adjacent arms); WDI—web depth as a percentage of longest arm length; WF—web formula (relative depth of web sectors, conventionally labelled A [between dorsal arm pair] to E [between ventral arm pair]). “I” following an abbreviation denotes an index, as a percentage of ML unless otherwise stipulated, and R or L denotes right or left.

Neither of the specimens redescribed in this contribution can be reliably assigned to a family within the superfamily Octopodoidea d’Orbigny, 1839, nor to a genus within any described family. Because of this, and for the purposes of the redescription that follows, the specimen identified as *Benthoctopus piscatorum* by Aldrich & Lu (1968) is herein referred to the genus “?*Benthoctopus*”; the reasons for this designation are discussed below. This specimen is also referred to as “cf. *piscatorum*”, to indicate that it resembles or is provisionally considered close to *piscatorum*, but the identification is not regarded as sufficiently certain to assign it unequivocally to this species. The type specimen of *Atlantoctopus pseudonymus* Grimpe is herein referred to *Vulcanoctopus*, although this generic assignment must likewise be considered uncertain. Alternative approaches include referring both specimens to *Vulcanoctopus* or designating them as “genus indeterminate”, but neither the author nor any reviewer of a previous iteration of this manuscript considered either option entirely satisfactory. This taxonomic ambiguity underscores a systemic challenge when working with historical type material for which molecular data are unavailable, rather than an isolated limitation of the current material. It will remain a problem until morphological characters, or character states, are identified that allow these genera to be unambiguously distinguished.

### SYSTEMATICS

Order Octopoda Leach, 1818

Suborder Incirrata Grimpe, 1916

Superfamily Octopodoidea d’Orbigny, 1839

Family *incertae sedis*

### Genus ?*Benthoctopus* Grimpe, 1921

If correctly assigned, the type species of this genus, by original designation, is *Octopus piscatorum* Verrill, 1879 (Grimpe, 1921: 299).

**Diagnosis (anatomical)**: octopuses with suckers in two rows; skin smooth; crop with diverticulum; ink sac absent.

?*Benthoctopus* cf. *piscatorum* (Verrill, 1879): Aldrich & Lu, 1968

?*Octopus piscatorum* Verrill, 1879: 470.

*Benthoctopus piscatorum* (Verrill)—Aldrich & Lu, 1968: 70–71, pl. 8 figs 1, 2; Nixon, 1991: 499–500; Nixon & Young, 2003: 321–322, figs 31.58, 59, 61, 62.

### Recognized distribution

Known from a single specimen, Placentia Bay, Newfoundland, 110 m depth.

### Material Examined

BM 1994037: ?*Benthoctopus* cf. *piscatorum*, female, ML *c*. 82 mm, 1 mile west of ‘Grass Island’, Placentia Bay, Newfoundland, 110 m, 28/10/1967, pres. M. Nixon.

### Description

Body incomplete, damaged; remnants of head and mantle separate from brachial crown (Fig. 1); funnel mostly free from ventral surface of head (FFuLI 19.5%), its length 31.8% ML (or 29.3% the ML of 89 mm reported by Aldrich & Lu (1968)). Remnants of mantle entirely smooth; originally purplish-brown (Aldrich & Lu, 1968: 70), now pallid. Webbing extends along the ventrolateral face of each arm almost to arm tip, and dorsolaterally to level of suckers 16–18 from the beak; maximum web depth (sectors A–C) 25% longest arm, and at its most shallow, 12% (sector E); WF A=B=C>D>E. Intact arms ∼56%–76% TL, 2.3–3.1× ML; both dorsolateral arms (arms II) damaged; their diameter where broken (relative to dorsal (I) and ventrolateral (III) and ventral (IV) arms) suggests a greater length, with an arm formula of I>II>III>IV. Intact arms with 68–80 suckers extending to arm tips; suckers moderately sized (ASI 5.1%–6.0%) (Table 1, 2). Gills with 8 inner and outer lamellae.

**Figure 1.**
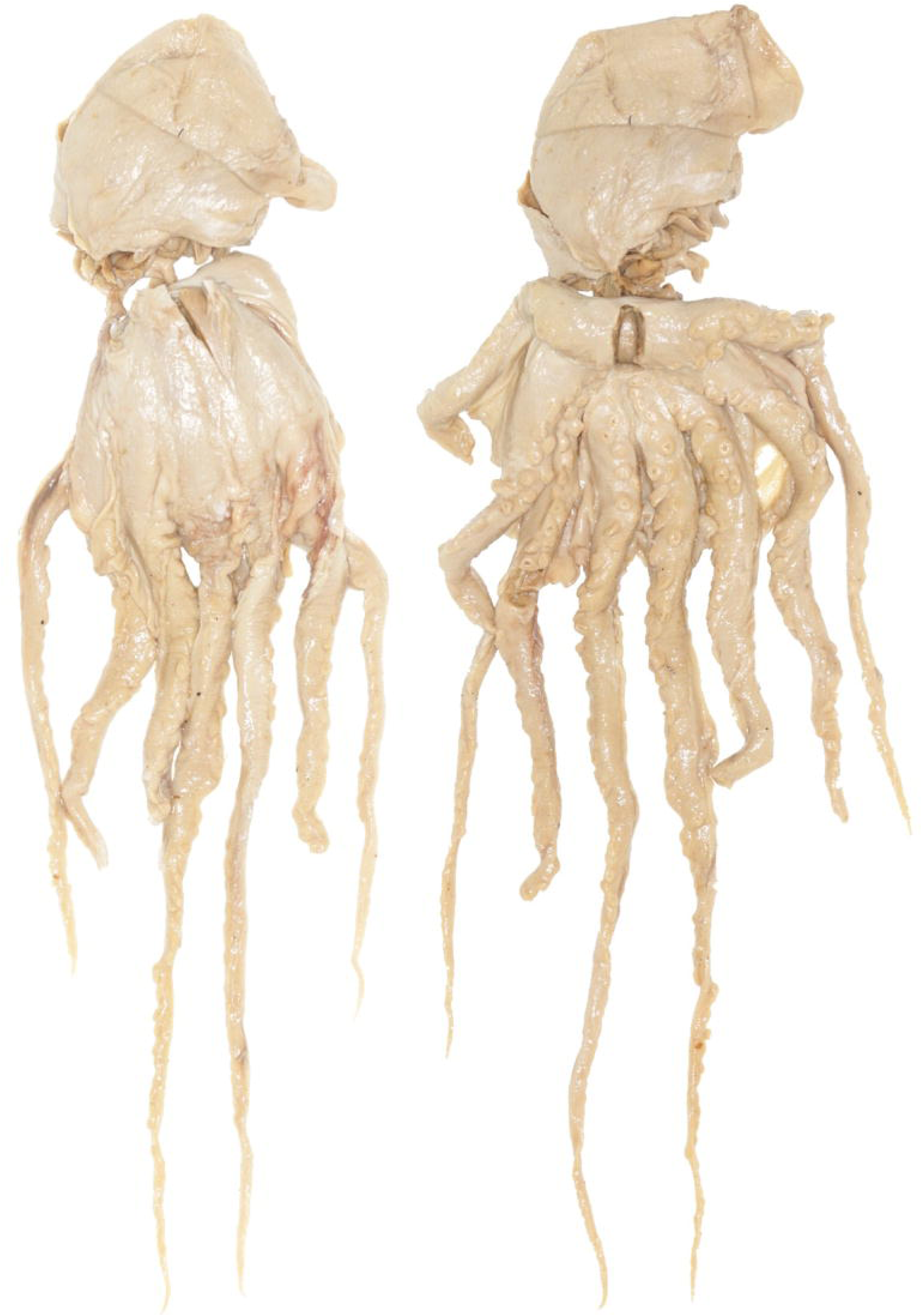
?*Benthoctopus* cf. *piscatorum* (Verrill) of Aldrich & Lu (1968): dorsal (left) and ventral (right) views (ML 82 mm).

**Table 1.** Raw measurements of *Benthoctopus piscatorum* (Verrill) holotype (USNM 574641; from O’Shea (1999: 193)); ?*Be.* cf. *piscatorum* (Verrill) of Aldrich & Lu (1968) (BM 1994037); *Be. pseudonymus* (Grimpe), holotype (MOM); and *Vulcanoctopus januarii* (Hoyle) lectotype (BMNH 1889.4.24.41).

|  | ***Be. piscatorum*** | **?*Be.* cf. *piscatorum*** | ***V. pseudonymus*** | ***V. januarii*** |
|---|---|---|---|---|
| TL | 150 | ~340[362] | 186 | 272 |
| ML | 40.0 | ~82[89] | 40.6 | 46 |
| MW | 34.5 | ~50 | 48.4 | 45 |
| HdL | 18.0 | - | 16.4 | 22 |
| HdW | 29.0 | - | 42.5 | 30 |
| EO | 5.0 | - | 3.2 | 1.9 |
| AL1R/L | 88/88 | 246/257 | 110*/119t | 152*/219* |
| AL2R/L | 74*/91 | 145*/130* | 132/133 | 198*/220 |
| AL3R/L | 83*/87 | 203/170* | 114*/121 | 133/217 |
| AL4R/L | 77/74 | 87*/192 | 118/114t | 162/175 |
| WDA | 30 | 63 | 38 | 34.4 |
| WDBR/L | */30 | 63/61 | 38/48 | 35.5*/42 |
| WDCR/L | */26 | 53/63 | 43/51 | 38.8*/31.9* |
| WDDR/L | */23 | 50/48 | 41/41 | 37.4/30.7* |
| WDE | 12* | 31 | 44 | 37 |
| ASC1R | 77 | 68/- | 87<sup>t</sup> | L135* |
| ASC2R | 64* | -/72 | 111 | L173 |
| ASC3R | 63* | -/- | 102 | 82/159 |
| ASC4R | 64 | -/80 | 106 | 155 |
| ASn1R | 2.5 | 4.9 | 2.5 | 3.7 |
| ASn2R | 2.0 | 4.2 | 2.5 | 3.7 |
| ASn3R/L | 2.5 | 4.2 | 2.5 | 3.4/3.2 |
| ASn4R | 2.5 | 4.2 | 2.5 | 3.2 |
| FuL | 11.0 | 26.1 | 16.7 | 19.4 |
| FFuL | 4.5 | ~16 | 4.4 | 5.4* |
| PA | - | - | 22 | 23.5 |
| GiLC | 7/7 | 8 | 6/6 | 9/9 |

Values in [brackets] are from Aldrich & Lu (1968); “-” indicates a missing value, usually because of damage. \* = damage, t = minor damage at extreme tip.

**Table 2.**
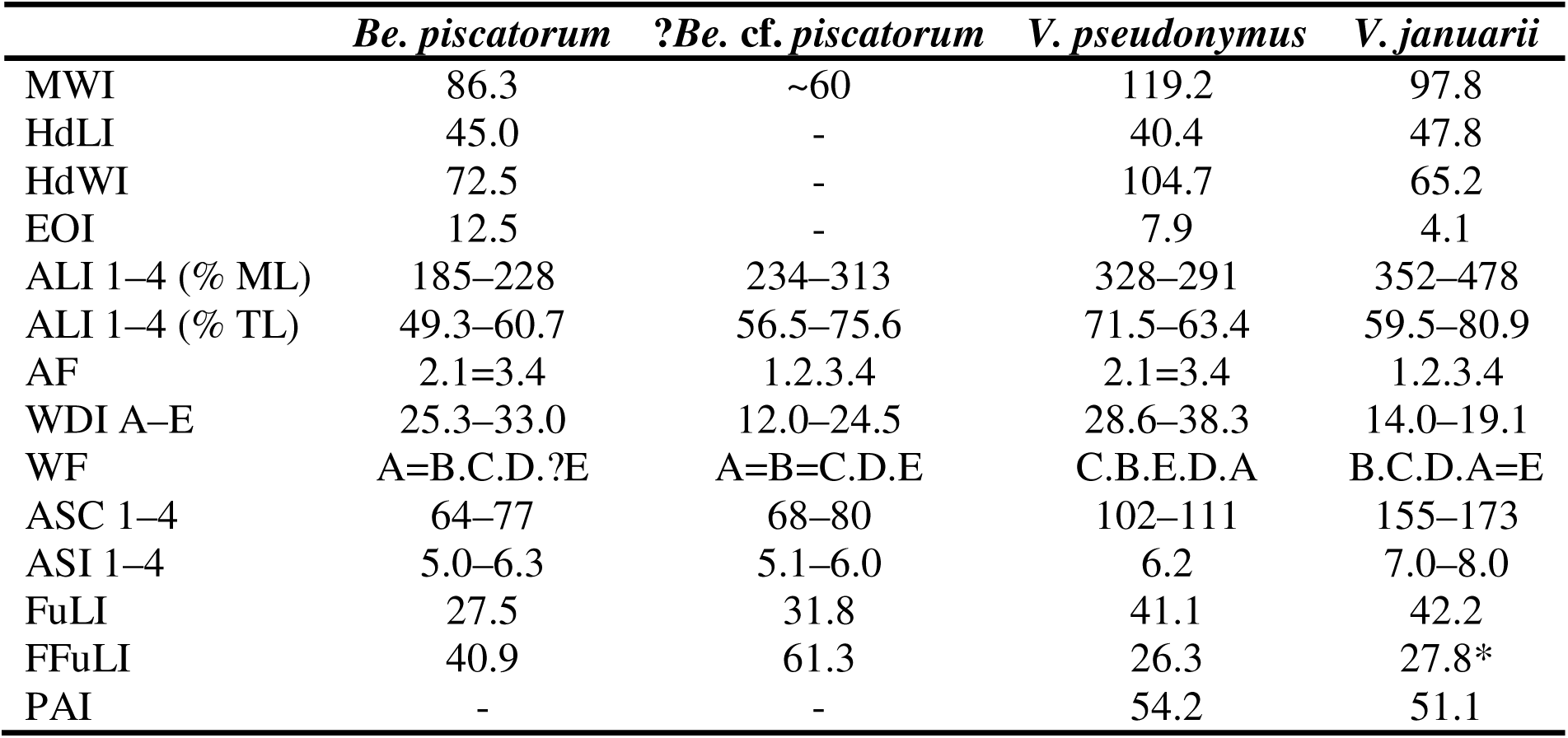
Select indices and counts of *Benthoctopus piscatorum* (Verrill) holotype (USNM 574641); ?*Be*. cf. *piscatorum* (Verrill) of Aldrich & Lu (1968) (BM 1994037); *Vulcanoctopus pseudonymus* (Grimpe) holotype (MOM); and *V. januarii* (Hoyle) lectotype (BMNH 1889.4.24.41).

|  | ***Be. piscatorum*** | **?*Be.* cf. *piscatorum*** | ***V. pseudonymus*** | ***V. januarii*** |
|---|---|---|---|---|
| MWI | 86.3 | ~60 | 119.2 | 97.8 |
| HdLI | 45.0 | - | 40.4 | 47.8 |
| HdWI | 72.5 | - | 104.7 | 65.2 |
| EOI | 12.5 | - | 7.9 | 4.1 |
| ALI 1–4 (% ML) | 185–228 | 234–313 | 328–291 | 352–478 |
| ALI 1–4 (% TL) | 49.3–60.7 | 56.5–75.6 | 71.5–63.4 | 59.5–80.9 |
| AF | 2.1=3.4 | 1.2.3.4 | 2.1=3.4 | 1.2.3.4 |
| WDI A–E | 25.3–33.0 | 12.0–24.5 | 28.6–38.3 | 14.0–19.1 |
| WF | A=B.C.D.?E | A=B=C.D.E | C.B.E.D.A | B.C.D.A=E |
| ASC 1–4 | 64–77 | 68–80 | 102–111 | 155–173 |
| ASI 1–4 | 5.0–6.3 | 5.1–6.0 | 6.2 | 7.0–8.0 |
| FuLI | 27.5 | 31.8 | 41.1 | 42.2 |
| FFuLI | 40.9 | 61.3 | 26.3 | 27.8* |
| PAI | - | - | 54.2 | 51.1 |

Alimentary canal (Fig. 2) with well-developed crop diverticulum (13% crop and distal esophagus length (to stomach)). BBL (length estimated from dimensions of cavity within arm crown, 17.8 mm along anteroposterior body axis) 20% ML (*fide* Aldrich & Lu, 1968) or 21.7% today. Posterior salivary glands (maximum dimension 18.2 mm) moderately sized (*sensu* O’Shea, 1999: 9, table 2), broadly triangular, 102% BBL, or 22.2% (based on current ML 82 mm) or 20% (based on original ML 89 mm) ML. Buccal bulb, radula, spiral caecum, and intestine presumed lost. Stomach tripartite, with thick muscular central section and thin-walled end sections, with non-compartmentalized lumen. Digestive gland (presumed) remnant with no vestige of ink sac, of dimensions (greatest) 27.1 mm × 17.7 mm.

**Figure 2.**
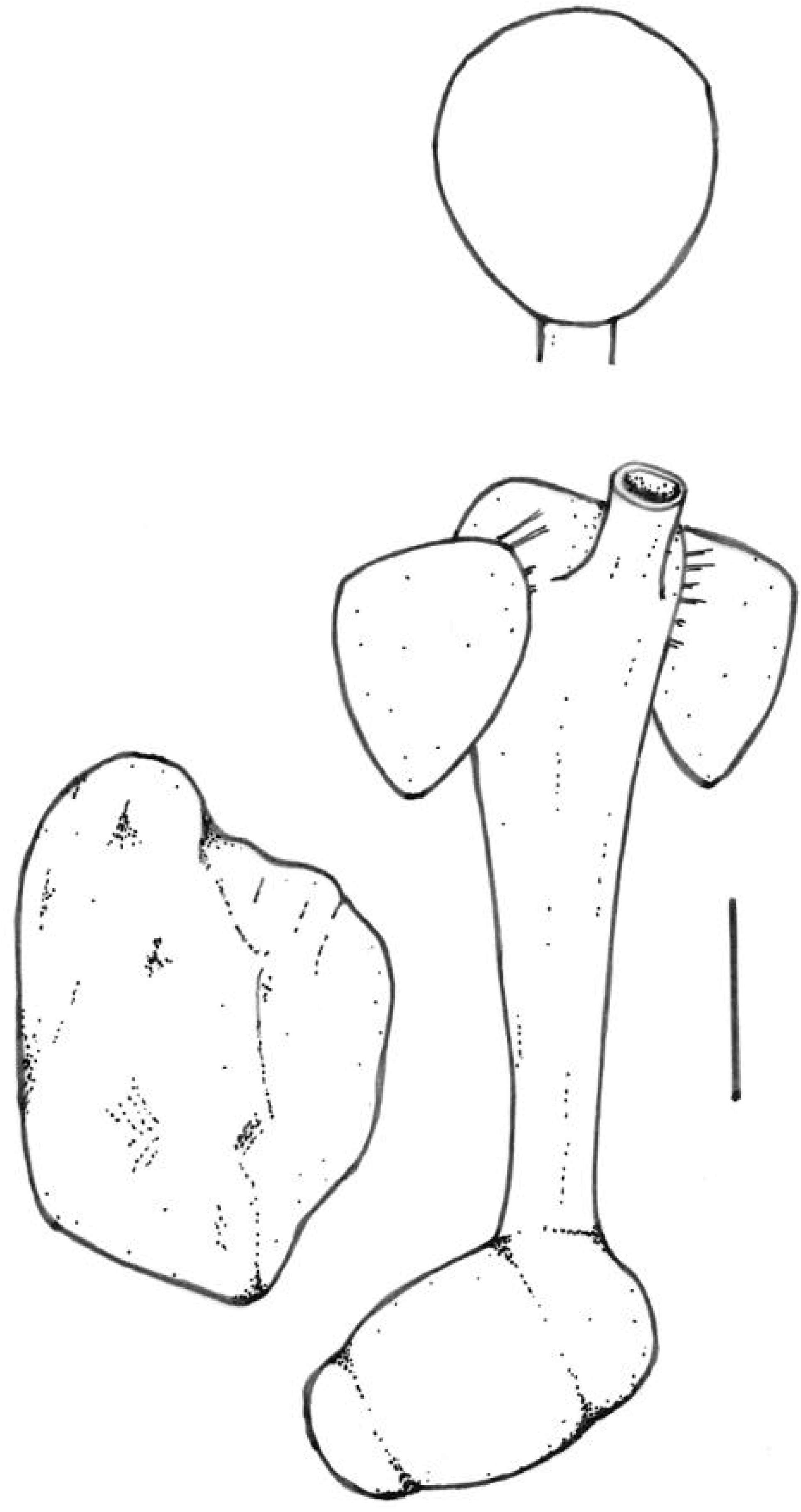
?*Benthoctopus* cf. *piscatorum* (Verrill) of Aldrich & Lu (1968): alimentary canal; scale bar 10 mm (the buccal bulb is lost; its size is estimated based on dimensions of the cavity between the arms). BB, buccal bulb; CD, crop diverticulum; DG, digestive gland (presumed); PSG, posterior salivary gland; ST, stomach.

Upper beak (Fig. 3, upper two images) moderately tall (height 93.1% length); hood and lateral walls dark brown, margins translucent; hood very deep (depth 52.2% beak length), with rounded crest, without posterior notch; rostrum triangular, tip slightly rounded; jaw straight, plunging near-vertically, without tooth-like prominence. Lower beak (Fig. 3, lower two images) depressed (height 66.7% width); hood, lateral wall, and wings dark brown, margins translucent. Lateral wall with deep basal notch and narrowly rounded crest. Hood deep (depth 37.8% beak length), with narrow triangular rostrum; hood with weak posterior notch; lateral wings very short (length 85.13% beak length).

**Figure 3.**
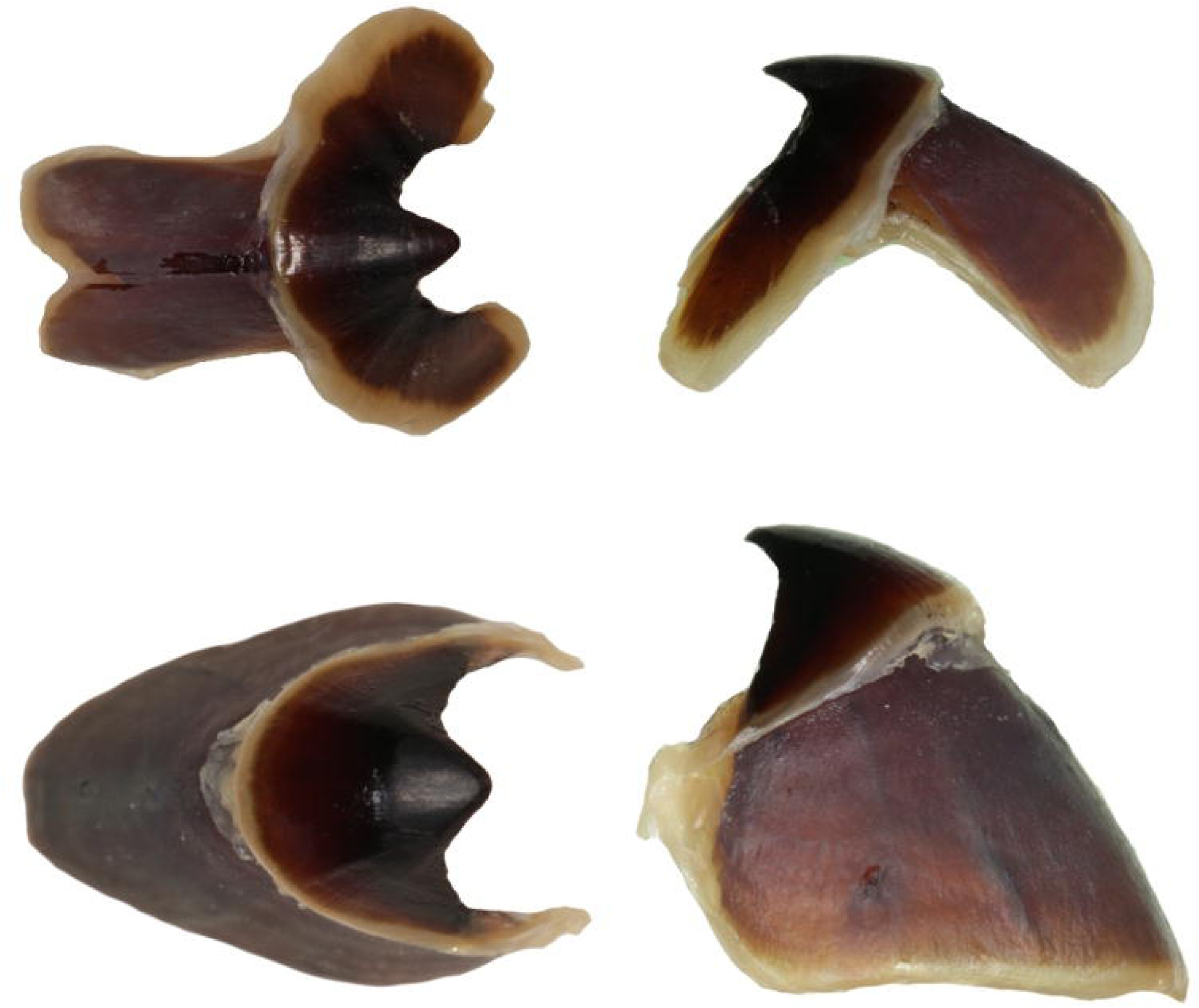
?*Benthoctopus* cf. *piscatorum* (Verrill) of Aldrich & Lu (1968): left (oral) and right (lateral) views of upper (upper two images), and lower (lower two images) beaks.

About half of ovary missing; loose mature ovarian eggs range 12.6–19.4 mm (16.3 ± 1.5 mm (standard deviation), n = 17) in length, and 4.6–5.8 mm (5.3 ± 0.3 mm, n = 15) in width. Oviductal gland (greatest dimension 13.9 mm) without apparent hemispheres, jet black; distal oviduct more swollen (proximal diameter 5.1 mm) than proximal duct (Fig. 4).

**Figure 4.**
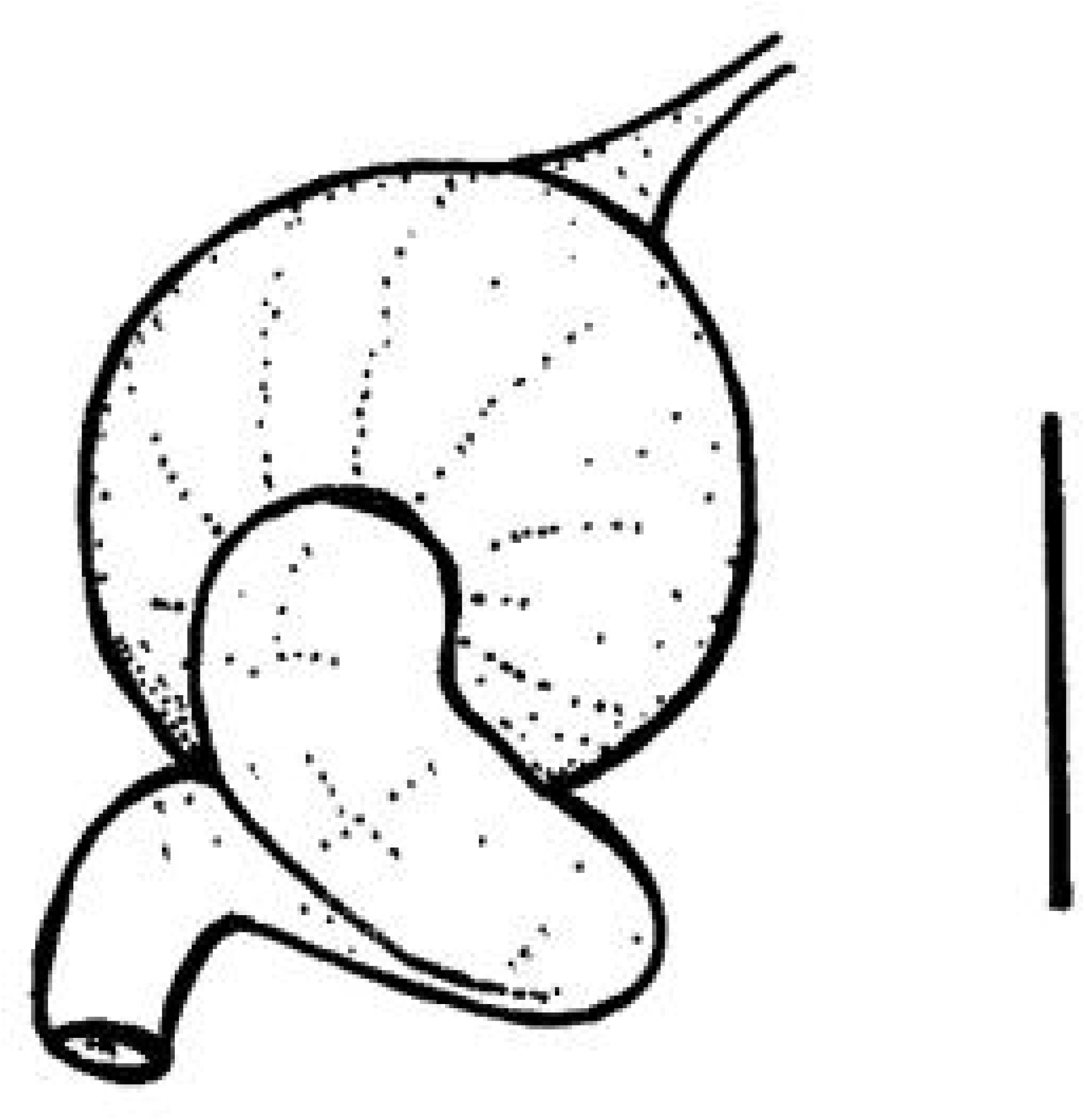
?*Benthoctopus* cf. *piscatorum* (Verrill) of Aldrich & Lu (1968): oviductal gland and oviduct; scale bar 10 mm.

### Remarks

In the absence of molecular data, candidate families to which this species likely belongs include Enteroctopodidae—for which member species have a very weakly or non-laminate ligula—or Bathypolypodidae—in which the ligula is strongly laminate. Because the specimen is female (as is the type of *Be. piscatorum*), no attempt is made to attribute this species to family (hence reference to *incertae sedis*). Consequently, the genus *Benthoctopus*, if this species is correctly assigned to it, is not diagnosed beyond characters discernable from this female specimen, that also have been routinely used in former generic diagnoses (ergo, no male characters are mentioned). This species is referred to *Benthoctopus* rather than *Vulcanoctopus* (*sensu* O’Shea et al., 2026) or *Bathypolypus* because it is more similar to the holotype of *Be. piscatorum* (the type species of this genus) than it is to any other named species in either genus, as detailed in the discussion. I reiterate, however, that I do not definitively attribute this specimen to Verrill’s species.

Although fragmentary and now in poor condition, the significance of this specimen does not depend upon its status as a name-bearing type, but rather upon the diagnostic morphological characters that remain available for examination. Dismissing it because it is no longer intact, or because it lacks type status—as recommended by several former reviewers of this manuscript—would overlook the information it continues to contribute to our understanding of the taxon.

### Enteroctopodidae Strugnell, Norman, Vecchione, Guzik & Allcock, 2014

*Vulcanoctopus* González & Guerra, 1998

**Diagnosis**: suckers in two rows; ink sac absent; ligula non-laminate.

“Vulcanoctopus” pseudonymus (Grimpe, 1922)

*Octopus levis* Hoyle, 1885—Joubin, 1900: 35, 36, pl. III fig. 5.

*Atlantoctopus pseudonymus* Grimpe, 1922: 41.

*Benthoctopus* (?) *pseudonymus* (Grimpe)—Robson, 1932: 248–249.

*Muusoctopus pseudonymus* (Grimpe)—Ibáñez et al., 2016: 6 (fig. 3).

Not *Octopus levis* Hoyle, 1885: 98, 99, pl. II figs 1–4, pl. III fig. 1.

### Recognized distribution

Known from the type locality only, west of the Azores, 1600 m.

### Material Examined

Holotype (Musée océanographique de Monaco): Princess Alice Stn 719, west of the Azores, 39°11’N, 32°44.5’W, 1600 m, 27 July 1896.

### Description

Female ML 40.6 mm, TL 186 mm (Fig. 5); MW 119.2% ML. Surfaces of mantle, head, arms, and web smooth, with subcutaneous gelatinous layer; ocular cirrus absent. Head length 40.4% ML, head width 104.7% ML; head scarcely demarcated from mantle (a string once tied between the head and brachial region exaggerates any demarcation). Orbits not meeting across dorsal midline of head; eye apertures dorsolaterally oriented; orifices puckered. Intact arms ∼63%–72% TL, 2.9–3.3× ML, with arms II longest, other arms of subequal length; AF II>I=III=IV. Web 29%–38% longest arm, with well-developed narrow membrane extending along ventrolateral arm face almost to arm tip (damaged on all arms); membrane extending along dorsolateral face of each arm ∼50%–75% arm length, to level of sucker 34 or 35; web formula B=C>E>D>A, deeper between left than right arm pairs. Suckers biserial to arm tips, small, ASIn 6.2 on all arms, none notably enlarged; intact total arm sucker counts 102–111.

**Figure 5.**
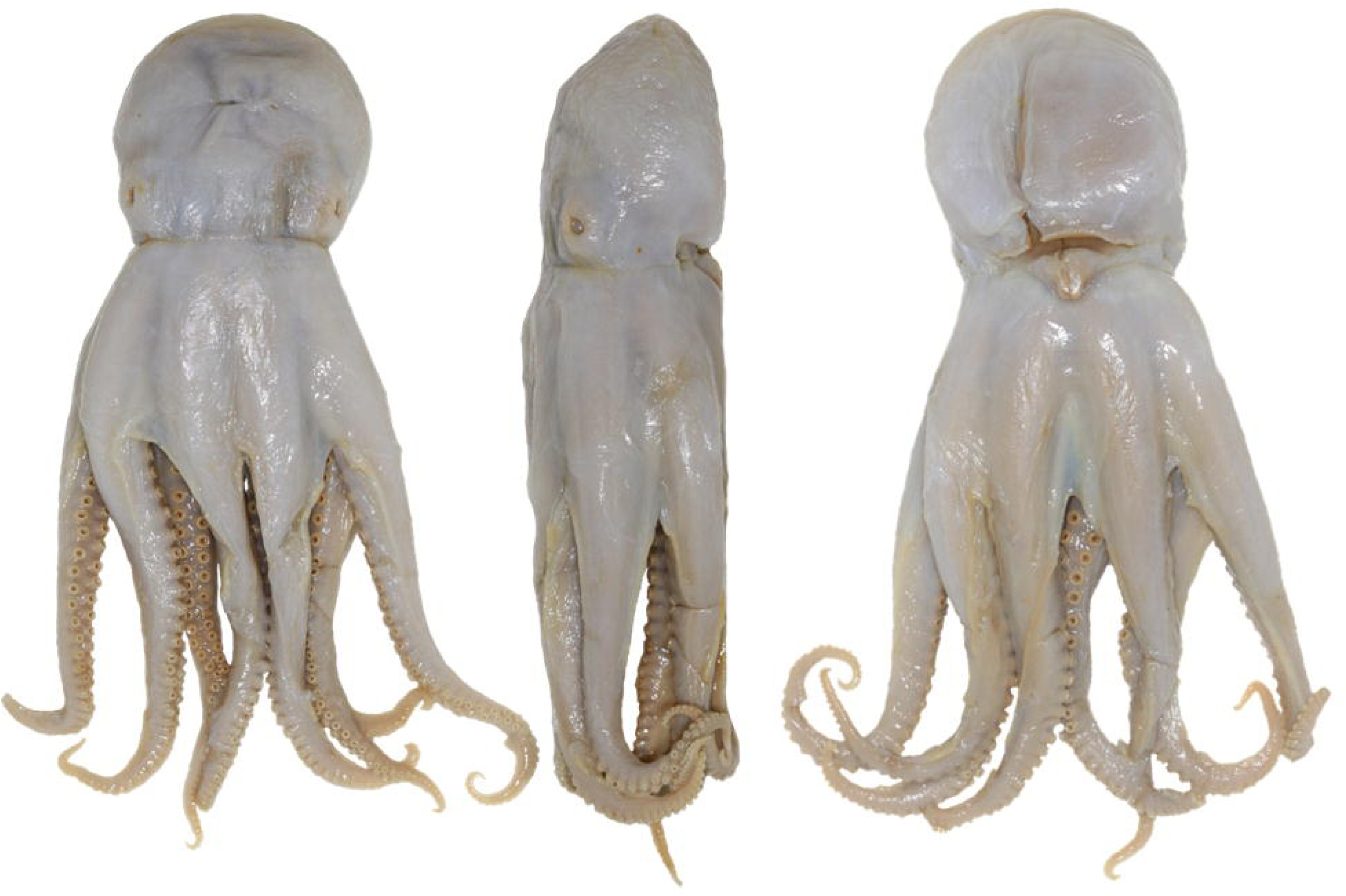
*Vulcanoctopus pseudonymus* (Grimpe, 1922): holotype (dorsal, lateral and ventral views; ML 40.6 mm).

Mantle with reduced pallial aperture (PAI 54.2%), transitional between types A and B of Robson (1926: 1334); FuLI 41.1% ML; adsiphonal pouches and funnel retractor muscles well developed; interpallial septum thick, extending along ventral, inner surface of mantle posteriorly to reproductive tissues, without connective membranes between renal tissues and inner, ventral surface of mantle cavity. Genital apertures opening lateral to and about half-way along septum; anal flaps absent. Gill length 28% ML (11.5 mm along efferent axis), with 6 inner and outer lamellae per demibranch; penultimate and terminal lamellae abruptly reduced. Chromatophores apparent on lateral muscles restraining digestive gland, but not renal papilla, anal region, or vestiges of visceral envelope. Funnel organ V V shaped, damaged, with comparably long inner and outer limbs.

Alimentary canal incomplete; crop with diverticulum; posterior salivary glands broadly heart-shaped, 22% ML (of moderate size, *sensu* O’Shea (1999: 9, table 2)) (based upon a loose gland within the visceral cavity). Female reproductive system represented by incomplete ovary sac, with paired oviducts, the proximal being appreciably narrower than the distal, with a beige-colored oviductal gland without apparent hemispheres.

### Remarks

For the same reasons detailed for Aldrich & Lu’s specimen, the genus and family to which Grimpe’s species belongs cannot be reliably determined, and confirmation of its generic and familial placement will require the collection and description of new material from or proximal to the type locality that is broadly consistent with the redescription of this taxon presented herein. That said, it is herein (with reservation) attributed to the genus *Vulcanoctopus*, and accordingly, family Enteroctopodidae, on grounds of gestalt.

The type of “*V*.” *pseudonymus* was initially described and measured after being fixed in formalin (après séjour dans le formol) (Joubin, 1900: 36)—a treatment that has rendered its tissues unsuitable for present-day molecular analyses, but which does explain the specimens’ excellent (for its age) condition over 120 years on. The lack of apparent chromatophores on the outer body surfaces is undoubtedly an artefact of long-term preservation, or treatment during this time, for the specimen was originally described as purplish “le corps est de couleur violacée, les membranes interbrachiales minces, légèrement translucides, sont de couleur un peu plus claire” (Joubin, 1900: 35). The puckered eye orifices suggest that it was fixed while alive, or in a premortem, relaxed condition, as does the presence of a well-developed, albeit now-damaged funnel organ. These attributes further suggest that the skin’s overall smoothness is real rather than an artefact, for animals capable of erecting papillae that have been fixed after plunging them live into preservatives are typically papillose (O’Shea, 1997). Joubin (1900: 35) also described the skin when fresh as completely smooth (“la peau est complètement lisse”).

Although the *pseudonymus* type remains relatively intact, it has been aggressively dissected and treated: the digestive gland is absent, the end of the crop diverticulum is severed, the impressions of a staple pierce the animal’s mantle dorsally, and a string has formerly constricted its brachial region. While Robson noted that this specimen lacked an ink sac, the nature of the incision is highly uncharacteristic of his dissection methods, suggesting the organ may have been removed or lost subsequent to his examination. Neither Joubin (who originally attributed this specimen to *Octopus levis*) nor Voss (who evidently examined this specimen in 1975, noted posthumously in Voss & Pearcy (1990: 48)) figured it or proffered any description of its anatomy; Robson (1932: 248) volunteered (without figures) that “the female reproductive organs resemble those of *B*[*athypolypus*]*. arcticus*, though (*a*) the oviduct is longer and (*b*) there is no second spermathecal swelling on the vagina”. It must be borne in mind that Muus (2002) regarded most of Robson’s *Ba. arcticus* to be *Ba. bairdii*. Robson’s reference to a second spermathecal swelling is most likely a reference to a dilatation of the distal oviduct, for he (Robson 1927: 259) referred to this as a “vaginal expansion” in *Benthoctopus sasaki* Robson, 1927 (now treated as a junior synonym of *Ba. arcticus*).

## DISCUSSION

### The Historical Ambiguity of *Octopus piscatorum*

On no occasion in which Voss referred to the type of *O. piscatorum* as being a species of *Bathypolypus* (Voss, 1988; Voss in Voss & Pearcy, 1990) did he indicate which species of *Bathypolypus* he thought it might be. At the time Voss (1988) diagnosed this genus as: “crop lacks diverticulum; radula lacks lateral cusps on rachidian; skin papillose; funnel organ VV; posterior salivary glands reduced”. Although these characters and their states are routinely used to differentiate species, they may have limited value for attributing a species to genus, or a genus to a family (Strugnell et al., 2014).

Muus (2002: 202) maintained that the senior synonym of *O. piscatorum* was *Ba. bairdii*—a synonymy subsequently recognized by Allcock et al. (2006). Further, Muus (2002: 185) alleged ‘I [Bent] discussed the *Bathypolypus*-*Benthoctopus* problems with Gilbert Voss [in 1981], who also examined the type specimen of *piscatorum* … [and who] … concurred with my opinion that Verrill (1879) described a somewhat atypical specimen of *Bathypolypus* and that, consequently, the genus *Benthoctopus* Grimpe, 1921 would be indistinguishable [from *Bathypolypus*]’.

### Morphological Integrity of the Placentia Bay Specimen

While in reasonably good condition (with a head) when figured by Nixon & Young (2003: fig. 31.58; the 5 mm scale bar therein being an obvious error for 5 cm), this specimen referred to herein as ?*Be*. cf. *piscatorum* is today extensively dissected and incomplete (Fig. 1). Nevertheless, important vestiges of its alimentary canal remain, as does the presumed digestive gland, within or upon which no trace of an ink sac is apparent (Fig. 2). Given the extensive experience of authors that have previously examined this specimen (Aldrich, Lu, Nixon, Young; see synonymy), and the apparent dissection of this mass, the presence of an ink sac would undoubtedly have been noted. Crucially, its well-developed crop diverticulum, smooth skin, and moderate-sized posterior salivary glands are fundamentally inconsistent with the historical diagnosis of *Bathypolypus* by Voss (1988).

### Divergence from Bathypolypus bairdii

Muus (2002) considered all octopuses attributed by Kumpf (1958) to *Ba. arcticus* (Prosch, 1847) from the Grand Banks, around Newfoundland, and from or near Placentia Bay, to be *Ba. bairdii*. Accordingly, if Aldrich & Lu’s specimen was misidentified, it would be most likely referable to *Ba. bairdii*. However, the Placentia Bay specimen differs demonstrably from *Ba. bairdii* (*sensu* Muus, 2002) across multiple characters (Table 3).

**Table 3.** Key diagnostic characters separating Aldrich & Lu’s Placentia Bay ?*Benthoctopus* cf. *piscatorum* from *Bathypolypus bairdii*.

| **Diagnostic Character** | **?*Benthoctopus* cf. *piscatorum* (Placentia Bay)** | ***Bathypolypus bairdii* (*sensu* Muus, 2002)** |
|---|---|---|
| Skin Sculpture | Entirely smooth | Densely papillose |
| Upper Beak Profile | Plunging jawline (Fig. 3) | Oblique jawline with a rounded, tooth-like prominence (Muus 2002: fig. 6b) |
| Total Length (TL) | ~340 mm (362 mm when fresh) | Rarely exceeds 200 mm |
| Mantle Length (ML) | ~82 mm today (89 mm fresh) | Maximum 75 mm (males); 68 mm (females) |
| Alimentary Tract | Well-developed crop diverticulum | Dilatation of the esophagus; no true diverticulum |
| Salivary Glands | Large; length 102% of estimated BBL | Reduced; length ~59.4% of BBL |

(Note: While ovarian egg length is variable, with Kumpf (1958: 53) citing lengths for *Ba. bairdii* to 12.8 mm and Muus (2002: 190) citing a range of 15–18 mm, the range of the preserved mature eggs of Aldrich & Lu’s specimen is negligibly greater at 12.6–19.4 mm [16.3 ± 1.5 mm, n = 17]).

Additionally, Aldrich & Lu’s specimen is physically unalike an exceptionally large specimen attributed to *Ba. bairdii* (ML c. 80 mm) from neighbouring waters (Saguenay Fjord, Québec) (Fig. 6). Ergo, if these aforementioned systematic characters are of any value—and there is no reason to believe otherwise—the Placentia Bay specimen must represent a second, sympatric inkless octopus species. The character states exhibited by the specimen are entirely inconsistent with those of *Ba. bairdii* (*sensu* Muus, 2002) but consistent with those historically attributed to *Benthoctopus* (*sensu* Robson, 1932). Because no other inkless species are currently recognized from this region, the question then becomes: to what species does it belong? By a process of elimination, it cannot be assigned to any currently accepted species, with the sole possible exception of *Be. piscatorum* (which is currently buried as a junior synonym of *Ba. bairdii*). Alternatively, it could represent a previously undescribed taxon or a named species currently treated as a junior synonym of another. In view of the latter possibility, it is compared with relevant taxa currently treated as synonyms of *Ba. arcticus* and *Ba. bairdii*.

**Figure 6.**
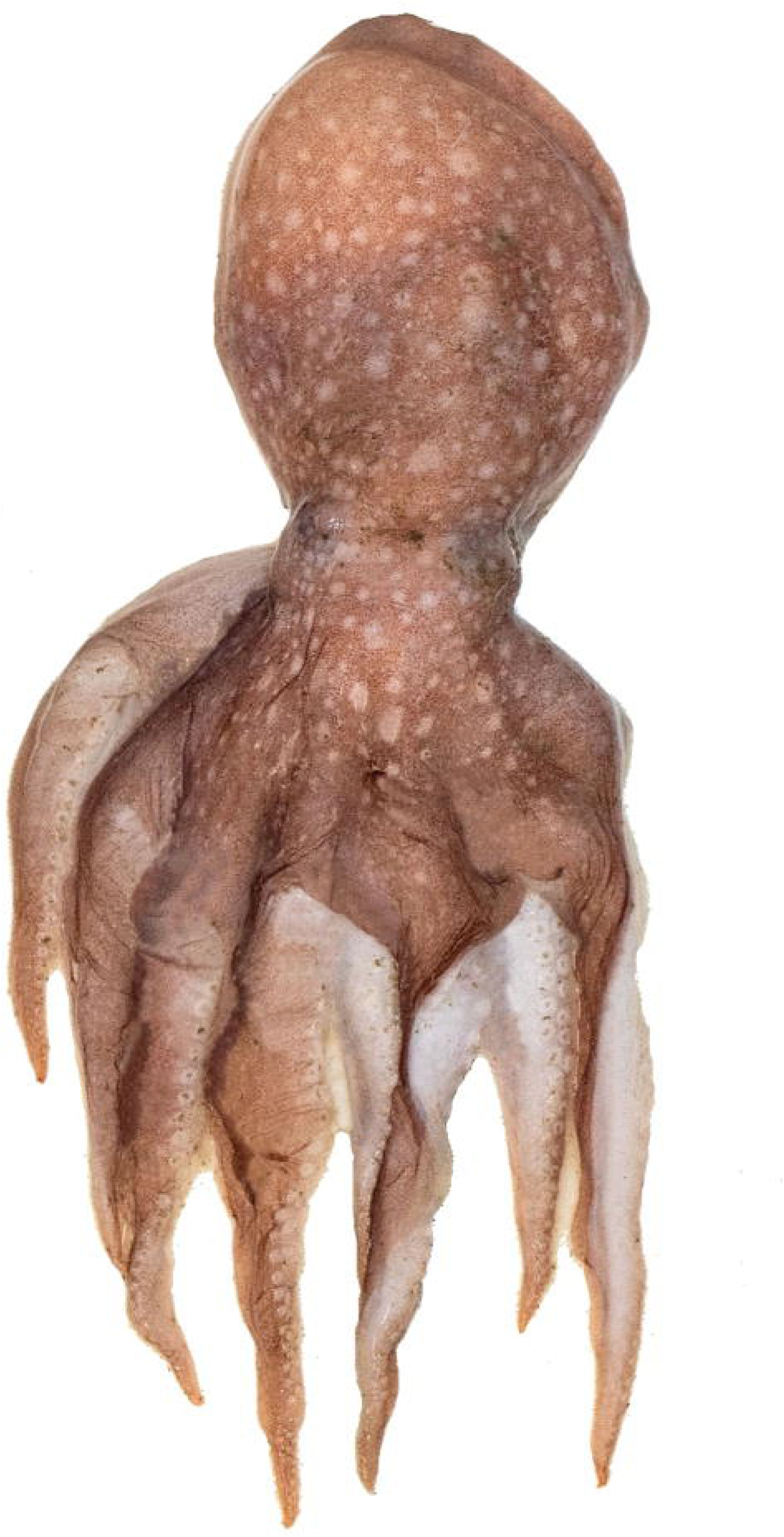
*Bathypolypus bairdii* (Verrill, 1873): dorsal view of fresh specimen from Saguenay Fjord, Québec (ML *c*. 80 mm). Photo courtesy of Claude Nozères, Fisheries and Oceans Canada.

### Unveiling Contradictions in the Current Nomenclatural Consensus

The justification for collapsing these taxa relies heavily on characters that exhibit untenable variation. Although Muus (2002: 184) referred to the damaged, partly missing funnel organ of the *O. piscatorum* type (which could be either a W or V V) as being of the “*Ba. bairdii* type”, the level of variation he reported for the latter (his fig. 18) included examples with separate U-shaped pads, completely fused W-shaped structures, and entirely separate linear limbs (IIII). Voss (1988) used this precise structural variation to differentiate *Benthoctopus* (W) from *Bathypolypus* (V V), and *Teretoctopus* (IIII). If Muus’s specimens were correctly identified, the shape of this structure might be less valuable for differentiating taxa than Voss (1977: 57) opined. Given its damage in the *O. piscatorum* holotype, the shape of this organ should not have been used to partly justify the synonymy of these two species. It should also be noted that Taite et al. (2023: 7) recently posited that *Ba. bairdii* may represent a species complex.

Muus maintained that Grimpe (1921) had been especially unlucky to have chosen *O. piscatorum* as the type species for *Benthoctopus*, claiming it was “identical with [his] *Ba. bairdii*” (Muus, 2002: 202). This assertion directly contradicts his earlier statement (p. 184) that this type was an “aberrant” individual, especially with regard to its unique SDEDI (sucker diameter as a percentage of eye diameter) index, which Muus used to demonstrate that the *Be. piscatorum* type had significantly larger suckers relative to eye size than any *Ba. bairdii*. Given this internal contradiction, and the completely smooth skin of the *O. piscatorum* type, Muus failed to unequivocally demonstrate that the type of *O. piscatorum* was conspecific with *Ba. bairdii*. While it remains possible that *O. piscatorum* represents a species of *Bathypolypus*, the formal synonymy of *Benthoctopus* with *Bathypolypus* based on historical data is indefensible.

### A Call for Nomenclatural Preservation

It follows that the generic name *Benthoctopus* and the species name *piscatorum* should be preserved as valid entities. They should remain distinct until the relationships between “*Be*.” *piscatorum*, the genus *Bathypolypus*, and *Ba. bairdii* can be independently and impartially corroborated. This reassessment must utilize expanded morphological datasets from respective type localities of each taxon and be fundamentally augmented with modern molecular data.

### Analytical Limitations Imposed by Historic Curation

The SDEDI index used by Muus (2002) to differentiate species cannot be calculated for Aldrich & Lu’s specimen, which now lacks a head and orbits, and no lenses remain; moreover, the resolution of figures in Aldrich & Lu (pl. 8, fig. 1) and Nixon & Young (2003: fig. 31.58) is too low to determine the original ratios of eye size to sucker diameter. Accordingly, this taxon can only be differentiated from others based on a more limited set of characters and states, most notably those associated with posterior salivary gland size relative to ML and BBL, the presence of a crop diverticulum, and for some taxa, beak morphology.

Unfortunately, key details regarding the alimentary canal are unavailable for the holotypes of *Be. piscatorum*, *“V.” pseudonymus*, and *V. januarii*. Further dissection of these specimens would permanently compromise their structural and cosmetic integrity, rendering them unsuitable for less-invasive modern examination by techniques such as computed tomography (CT).

### Comparisons within the Genus *Bathypolypus*

Of other North Atlantic inkless octopuses, *Ba. arcticus* represents the second-most-likely name to apply to Aldrich & Lu’s specimen had it been misidentified. However, the recognized southern limit of *Ba. arcticus* (∼60°N, *fide* Muus, 2002: fig. 20) lies well north of Placentia Bay (∼47°N), and the species is typically extensively papillose. Furthermore, based on historical anatomical illustrations (*fide* Muus, 2002: 180, fig. 2), its crop diverticulum spans almost 30% of the combined length of the crop and distal esophagus, the buccal bulb is small (∼20% ML), and the posterior salivary glands are vestigial at 14% ML (72.6% BBL, *sensu* O’Shea, 1999: 9, table 2). While both *Ba. arcticus* and ?*Be*. cf. *piscatorum* have a well-developed crop diverticulum, their other anatomical proportions differ. Combined with a discontinuous geographic distribution, it is highly improbable that the Placentia Bay specimen is conspecific with *Ba. arcticus*.

A third species, *Ba. pugniger* Muus, 2002, has posterior salivary glands spanning 23% ML—comparable to the Placentia Bay specimen—but differs sharply in being extensively papillose, significantly smaller (maximum ML of 54 mm across 31 examined specimens), lacking a crop diverticulum, and having a non-plunging upper beak jawline with a rounded, tooth-like prominence.

A fourth species, *Ba. faeroensis* Russell, 1909 (*sensu* Toll, 1985), warrants comparison also. Muus (2002: 184) synonymized *Ba. faeroensis* with *Ba. arcticus*, alleging that the two were indistinguishable “in all aspects”. This assertion is patently incorrect: *Bathypolypus faeroensis* is larger (ML 74.5 mm), whereas the mantle of *Ba*. *arcticus* rarely exceeds 70 mm (Muus, 2002: 178); it has lower arm sucker counts (70–79 on Toll’s neotype of *Ba. faeroensis* (O’Shea, 1999: 194) and between 60 and 70 for a large syntype female (Russell, 1909: 448) vs. 80–90 on the dorsal arms and 60–70 on the ventral arms for *Ba*. *arcticus* (Muus, 2002)); and gill lamellae counts differ (9 or 10 outer lamellae (Toll, 1985), compared to “6–7(8)” [sic] (Muus, 2002: 178)). Unfortunately, Toll (1985) neither mentioned nor illustrated the posterior salivary glands of *Ba. faeroensis*, and I, when I last examined this specimen in the mid-1990s, and figured it (O’Shea, 1999: fig. 115C, D), failed to note any detail also. Until compelling evidence demonstrates otherwise, *Ba. faeroensis* should be treated as a distinct, valid taxon.

### Intergeneric Separation from *Vulcanoctopus* Taxa

From *Vulcanoctopus* taxa—to which both ?*Be.* cf. *piscatorum* and possibly the type specimen of *Be. piscatorum* (Verrill), and “*V.*” *pseudonymus* plausibly belong—Aldrich and Lu’s specimen differs from the two other North Atlantic species recognized from this region by Ibáñez et al. (2016): *V. januarii* and *V. johnsonianus* (Allcock, Strugnell, Ruggiero & Collins, 2006) comb. nov., and from *V. normani* (Massy, 1907) comb. nov. also—a third species considered by Gleadall (2013) to be a junior synonym of *V. januarii*. While Massy (1909) first raised the possibility that *V. normani* was a synonym of *Be. piscatorum* when she described it, and Robson (1932) later effected the synonymy, Allcock et al. (2006) resurrected and redescribed this species, and, given the holotype had been lost, designated a lectotype for it. In the following discussion, the rationale of Taite et al. (2023) is followed and Massy’s *normani* (*sensu* Allcock et al., 2006) is treated as distinct from Hoyle’s *januarii*.

To help clarify the dense morphological boundaries separating ?*Be*. cf. *piscatorum* from the recognized North Atlantic *Vulcanoctopus* species—*V. januarii* (Hoyle, 1885), *V. normani* (Massy, 1907), and *V. johnsonianus* (Allcock et al., 2006)—the key diagnostic metrics are detailed in Table 4.

**Table 4.** Key diagnostic characters separating Aldrich & Lu’s Placentia Bay ?*Benthoctopus* cf. *piscatorum* from North Atlantic *Vulcanoctopus* taxa.

| **Diagnostic Character** | **?*Be.* cf. *piscatorum*<br>(Placentia Bay)** | ***V. januarii*** | ***V. normani*** | ***V. johnsonianus*** |
|---|---|---|---|---|
| Total (intact) Arm Sucker Count (ASC) | 68–80 | 155–173<sup>a</sup> | 144–199 | 130–145 |
| Normal Sucker Index (ASIn) | 5.1%–5.9% | 7.0%–8.0%<sup>a</sup> | 5.0%–6.0% | Data variable |
| Crop Diverticulum | Well-developed | Absent<sup>b</sup> | Absent | Absent |
| Posterior Salivary Gland / BBL | 102% | Undetermined | 66.8% | 68.2% |
| Buccal Bulb Length (BBL) / ML | 20%–21.7%<sup>c</sup> | Undetermined | 38.8% | Data variable |
| Upper Beak Jawline | Plunging | Oblique | Oblique / Rounded | Plunging |
| Gill Lamellae Count (GiLC) | 8 | 9 | 7–10 | 8–11 |

(Note: <sup>a</sup>, lectotype; <sup>b</sup>, from Toll (1981); <sup>c</sup> 20% ML (*fide* Aldrich & Lu, 1968) or 21.7% (today))

Because Massy’s *normani* is treated as a valid taxon, those figures provided for it by Allcock et al. (2006) cannot be used as a proxy for the anatomy of *januarii* (whereas Gleadall regarded the molecular sequence of *normani* to be a proxy for that of *januarii*). However, Gleadall (2004) also included in his synonymy of *januarii* those specimens attributed to this species by Toll (1981), who also described and illustrated this species’ anatomy. The most pronounced divergence between the Placentia Bay specimen and *V. januarii* proves to be in total arm sucker count (68–80 vs. 155–173). However, as illustrated by Toll (1981), *V. januarii* also differs fundamentally in lacking a crop diverticulum and possessing an oblique upper beak jawline. Until new material is collected from the type locality off Barra Grande, Brazil, and redescribed, *V. januarii* will remain poorly known, and figures for *V. normani* should not be used as a proxy for its anatomy.

The divergence from *V. normani* is equally pronounced. The considerable differences in ASC between comparably sized individuals make it highly improbable that they are conspecific. Furthermore, *V. normani* has a non-plunging, distinctly different beak morphology (Allcock et al., 2006: fig. 3A), lacks a crop diverticulum, and has a much larger buccal bulb relative to mantle length (BBL 38.8% ML vs. 20%–21.7% ML in ?*Be*. cf. *piscatorum*) (Allcock et al., 2006: fig. 3B).

While ?*Be*. cf. *piscatorum* and *V. johnsonianus* differ fundamentally in their posterior salivary-to-buccal mass proportions (102% vs. 68.2%), relative web depths (12.0%–24.5% vs. 16.0%–30.0%), and total (intact) arm sucker counts, they are similar in several characters. Both taxa have highly similar upper and lower beak morphologies—specifically the upper beak’s straight, plunging jawline, and they have a similar crop-to-esophagus ratio (13% vs. 18%). In these characteristics these two taxa are more similar to each other than either is to *V. normani*.

It is noteworthy that each North Atlantic *Vulcanoctopus* taxon lacks a crop diverticulum, but the presence of this diverticulum was formerly considered to be diagnostic of the genus *Benthoctopus* (to which each of these taxa has been historically referred).

### Comparative Integrity of “*Vulcanoctopus*” *pseudonymus*

There are no remarkable morphological differences between the type of *Be. piscatorum*, Aldrich & Lu’s ?*Be.* cf. *piscatorum*, and *V. pseudonymus*; each is smooth bodied, female, and characterized by a relatively low total arm sucker count. However, the type of *Be. piscatorum* and the Placentia Bay specimen are distinctly muscular-bodied, whereas “*V.*” *pseudonymus* is more gelatinous. Furthermore, these three taxa differ in gill lamellae counts, which are lowest for “*V.*” *pseudonymus* (6), intermediate in the *piscatorum* type and Placentia Bay specimen (7 and 8 respectively), and highest for *V. januarii* (9)). These distinctions indicate that neither *Be. piscatorum* nor ?*Be*. cf. *piscatorum* are likely to be conspecific with “*V.*” *pseudonymus*.

Rather than forcing “*V*.” *pseudonymus* into *Benthoctopus*, it is provisionally referred here to *Vulcanoctopus*. Unfortunately, the type specimens of *V. hydrothermalis* González & Guerra, 1998 cannot be located in the collections of the Muséum national d’Histoire naturelle, despite extensive physical searches by the author. Because no recent loan documentation exists, these type materials are presently categorized as “misplaced” rather than lost. Nevertheless, published data show that *V. hydrothermalis* and “*V*.” *pseudonymus* differ markedly at comparable mantle lengths. Arm sucker counts of *V. hydrothermalis* are highly variable, ranging 38–87 per sucker *row* along each arm (= 76–174 suckers per arm: Gonzalez et al. (1998, 2002)), or 90–126 per *half arm* (Gonzalez et al., 2008) for a single female. Problematically, from a taxonomic perspective, they also vary geographically (Voight, 2012) and ontogenetically (Gonzalez et al., 2002). However, in addition to “*V.*” *pseudonymus* having fewer suckers (102–111 in total) it has 6 lamellae per gill demibranch, compared to 7 or (primarily) 8 in *V. hydrothermalis*.

While no active hydrothermal vents are recorded from the reported collection site of “*V*.” *pseudonymus*, vent fields are well documented across the Azores region, with the Menez Gwen field on the Mid-Atlantic Ridge representing the closest viable habitat proxy (Sarrazin et al., 2022).

### Historical Systematics and *Bathypolypus ergasticus*

Robson (1932: 231) considered *Be. pseudonymus* to be a possible synonym of *Atlantoctopus lothei* (Chun, 1913), which he also considered to be a possible synonym of *Ba. ergasticus* (Fischer & Fischer, 1892). Any affinity to *Ba. ergasticus* can be definitively dismissed here; its extraordinarily high arm sucker count (> 200) separates it from “*V.*” *pseudonymus* (Guerra et al., 2014: 617; pers obsv.).

### Synthesis and Taxonomic Resolution

By a systematic process of elimination, Aldrich & Lu’s Placentia Bay specimen remains distinct from all currently accepted *Bathypolypus* and *Vulcanoctopus* taxa in the North Atlantic, and from various species that have been considered to be synonyms of a number of these species. Unless it represents an entirely un-described species, it is morphologically most similar to—and possibly conspecific with—the holotype of *Be. piscatorum*. Because both historical specimens are solitary, damaged females, a definitive union remains limited by the absence of male hectocotylus data. However, dismissing the Placentia Bay specimen as “aberrant” (as Muus did with the *Be. piscatorum* holotype) or disregarding it due to its preservation state is a highly inappropriate systematic course of action.

To resolve these deep nomenclatural conflicts and map out historical priority, future taxonomic work should follow a framework that involves: 1) collection of topotypic material from respective type localities; 2) integration of morphological and molecular data; and 3) resolution of the identity of *Teretoctopus* Robson, 1929 (*sensu* O’Shea et al., 2026).

## ACKNOWLEDGEMENTS

I wish to thank Claude Nozères and Johanne Gauthier, Fisheries and Oceans Canada, for valued discussions on octopuses off Newfoundland, and an image of one specimen reproduced herein from Saguenay Fjord, Québec. Dr Chung-Cheng Lu is also thanked for discussions regarding the specimen that he described with the late Dr Frederick Aldrich many years earlier. Thanks are extended to Dr Amanda (Mandy) Reid, at the time at the Australian Museum (Sydney, Australia), for officiating the loan of material referred to herein, providing facilities for the author while he researched there, and for being a very good friend. Special thanks also go to Jonathan Ablett of the Natural History Museum, London, UK, and Michèle Bruni, Musée océanographique de Monaco, Monaco, France, for the loan of this material. Finally, I thank the reviewers whose constructive feedback during earlier development significantly helped refine and improve the precision of this text.

## Conflict of interest

The author declares no conflict of interest.

## Ethical approval

No animal testing was performed during this study (materials referred to herein have been accessioned into museum collections).

## Sampling and field studies

The study is based on historical museum-preserved specimens, for which it can only be assumed that any (if appropriate) permit was used during specimen collection.

## Data availability

All data generated or analyzed during this study are included in this published article.

## Author contribution

The sole author is responsible for all aspects of this manuscript.

## Notes

### Competing Interest Statement

The authors have declared no competing interest.

### Summary of Updates

This revised version incorporates amendments to the manuscript, including clarification of the morphological evidence, further consideration of the historical taxonomy and nomenclature, corrections to the text and references, and line and figure numbers. The central subject and objectives of the contribution remain unchanged.

